# Topological Assembly of a Deployable Hoberman Flight Ring from DNA

**DOI:** 10.1101/2020.09.09.290197

**Authors:** Ruixin Li, Haorong Chen, Jong Hyun Choi

**Affiliations:** School of Mechanical Engineering, Purdue University, West Lafayette, Indiana 47907

**Keywords:** DNA origami, self-assembly, topology, deployable, auxetic, reconfiguration

## Abstract

Deployable geometries are finite auxetic structures that preserve their overall shapes during expansion and contraction. The topological behaviors emerge from intricately arranged elements and their connections. Despite considerable utility of such configurations in nature and in engineering, deployable nanostructures have never been demonstrated. Here we show a deployable flight ring, a simplified planar structure of Hoberman sphere, using DNA origami. The DNA flight ring consists of topologically assembled six triangles in two layers that can slide against each other, thereby switching between two distinct (open and closed) states. The origami topology is a trefoil knot, and its auxetic reconfiguration results in negative Poisson’s ratios. This work shows the feasibility of deployable nanoarchitectures, providing a versatile platform for topological studies and opening new opportunities for bioengineering.

---

Deployable geometries are two-dimensional (2D) or three-dimensional (3D) finite structures that preserve their global shapes during expansion and contraction.^1^ The structural transformation shows auxetic behaviors, which may be characterized by a negative Poisson’s ratio (*ν*). Such topological behaviors emerge from their unique, intricate geometrical designs. In a Jitterbug transformer, for example, eight rigid equilateral triangles are linked at vertices, and the triangles can rotate around the linkages.^2^ This allows deployable reconfigurations between octahedron (one of five Platonic solids) and cuboctahedron (one of thirteen Archimedean solids). By varying the shape and length of elements (regular polygons) in the transformer, multiple variants such as other deployable Platonic and Archimedean solids have been studied mathematically.^3^ Another example is the Hoberman sphere, a commercially available, popular toy for kids and a possible choice for sculptures. The Hoberman sphere is also formed by rigid edges with flexible linkages,^4^ and can shrink and expand by scissor-like actions at the joints while maintaining the overall spherical shape. In theory, it can be constructed regardless of materials and lengthscale. There is a 6-m-diameter Hoberman sphere from aluminum in the AHHAA Science Center in Tartu, Estonia, while the commercial toys made of plastics are typically about 10 cm in length.^5^ In nature, the shell of cowpea chlorotic mottle virus (CCMV) is also considered as a deployable structure.^6,7^ Despite the considerable utility of deployable configurations in nature and in engineering, synthetic deployable nanostructures have not been available to date, to the best of our knowledge.

Here we demonstrate a deployable nanoscale flight ring, inspired by the Hoberman sphere, using DNA origami. As a bottom-up biomolecular self-assembly,^8–10^ DNA origami is a powerful method to create topological and dynamic structures of arbitrary shapes with excellent programmability and structural predictability.^11–19^ While the 3D Hoberman sphere is originally constructed by rigid rods linked at both ends and in the middle, its planar structure, flight ring, has two layers of interconnected rods.^20^ In this work, we design the 2D deployable structure by combining the rods into six equilateral triangles arranged in two layers. The triangles are made of a topologically routed scaffold strand stabilized by a set of staple strands, together forming a trefoil knot. The nanoscale DNA flight ring can switch between two distinct, open and closed states by sliding the triangles against each other via strand displacement.^21,22^ The deployable reconfiguration shows the features of both Jitterbug transformers and Hoberman structures: (i) equilateral triangles as elements and flexible linkages at the vertices, and (ii) two slidable layers. The origami structures and their dynamic reconfigurations are studied with atomic force microscopy (AFM). The auxetic deformation is observed with the Poisson’s ratio ranging from approximately −1.67 to −2.0.

Figure 1a and b show a Hoberman sphere toy and its planar structure, respectively. The basic mechanism in the auxetic structural transformation is hinging of the rings made of stiff rods in two layers linked with scissor-like interlayer joints^23^. Our first step of simplification is a reduction of the number of rods such that the two-layered rings are represented by two equilateral triangles (Figure 1c), where the red triangle is on top of the blue triangle. As the hinged parts (where the blue and red lines cross) are pushed in, the inscribed (dashed) circle shrinks while the circumscribed (dashed) circle maintains its size (Figure 1d). If the hinges are pushed back, the process is reversed, restoring the original configuration in Figure 1c. Since it is a challenge to construct multiple scissor-like joints from DNA in a single structure, we further simplify the planar geometry with two sets of triangles (red triangles on top of blue triangles). The hinging mechanism with scissor-like joints is replaced by sliding of the triangles in two layers, as shown in Figure 1e and f. Three inside vertices are designed, and each connects two triangles in different color (*i.e.*, different layer) on the opposite side. The sliding of the triangles around the linkages will lead to two distinct states. When the red triangles are located perfectly on top of the blue triangles, the inscribed (dashed) circle will be largest, that is, the open state as depicted in Figure 1e. In contrast, the inscribed (dashed) circle becomes smallest if the triangles slide the farthest around the interlayer linkages, forming a regular hexagon (*i.e.*, the closed state as illustrated in Figure 1f).

**Figure 1.**
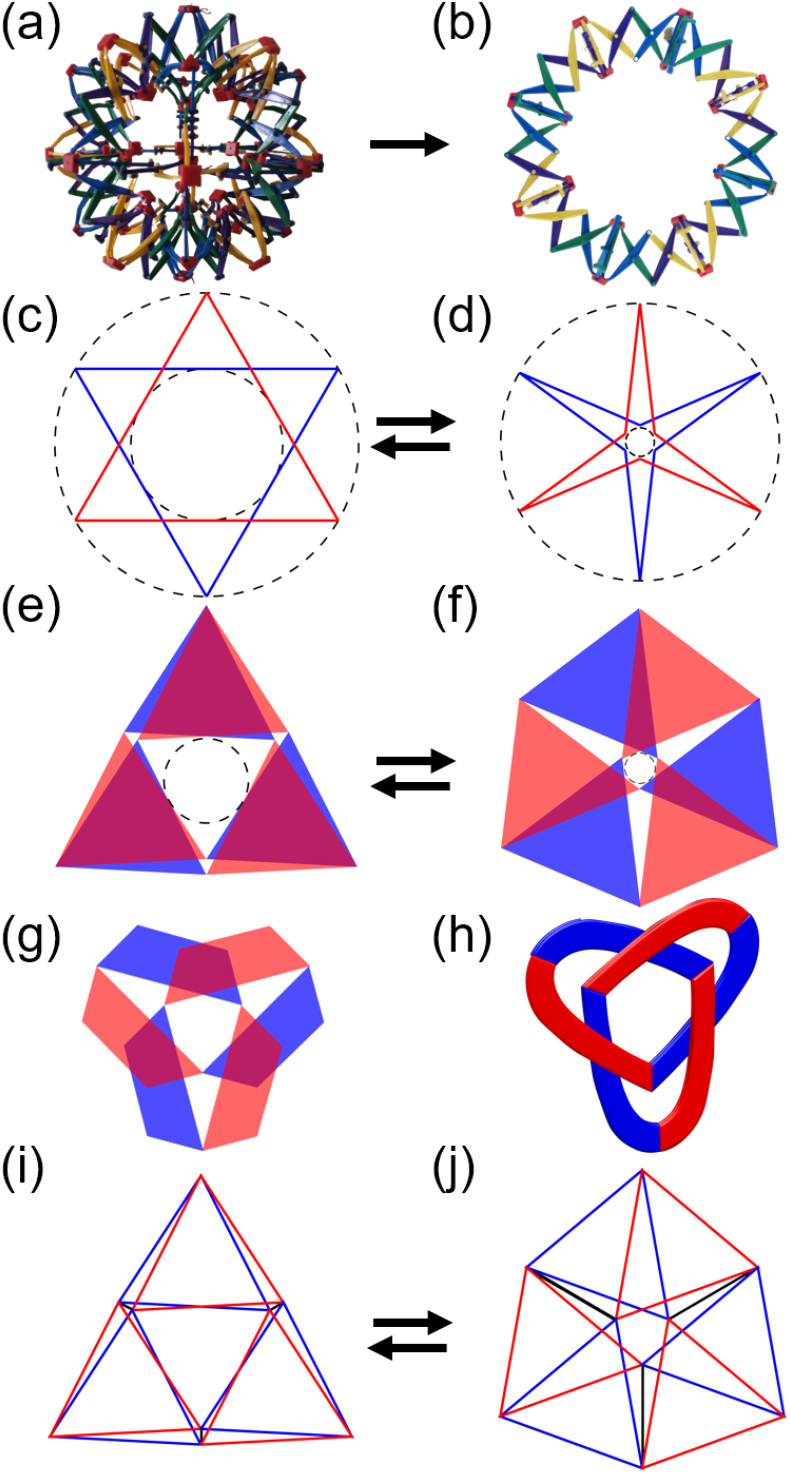
DNA origami flight ring. (a) 3D Hoberman sphere. (b) 2D Planar structure of Hoberman sphere. (c) – (d) Conceptual design of a flight ring in two distinct states with two (red and blue) layers. Similar to (b), the two layers slide against each other around the interlayer joints in the middle of edges. The movement of the two layers can lead to dynamic switching between open (c) and closed (d) states. (e) – (f) Intermediate design of the flight ring in open (e) and closed (f) states. The three red triangles are on top of the blue triangles. Note that the inside vertex of each red triangle is connected to that of the blue triangle on the opposite side. (g) Geometrical representation of the structures in (e) and (f), which is a trefoil knot (h). (i) – (j) Final design of the DNA flight ring. Wireframe DNA origami with red and blue edges is used rather than solid triangles. By modulating the length of jack edges (shown in black), the flight ring assumes the open (i) or closed (j) conformation.

The deployable configuration in our design is a trefoil knot.^24^ Consider the triangles are reduced to the bands with the identical color scheme (Figure 1g). With the red band going over the blue band in Figure 1h, this flight ring can be seen as a left-handed trefoil knot (Figure S2). This implies that as the flight ring is built by routing a circular DNA scaffold, the strand is folded into a trefoil knot, which is not a violation to the laws of topology. For example, a circular single-stranded DNA (ssDNA) may be transformed into a trefoil knot by having staples bring parts together to form a circular double-stranded DNA (dsDNA). In the actual design, we realize the triangular subunits using wireframe DNA origami^25,26^ (Figure 1i, j), where the three edges in each subunit are constructed with two dsDNA bundles. All the joints are made of unpaired ssDNA segments of scaffold with various lengths depending on whether or not the vertex rotates. For example, interlayer joints rotate, while intralayer joints do not. Thus, the scaffold strand goes through all the edges and joints (Figure S3). Since the edges are formed by two-dsDNA bundles, each triangle is routed by the scaffold twice in the antiparallel directions. Staples are placed to stabilize the folded scaffold pattern, thus forming the edges and joints.

To enable deployable reconfiguration, we implement two-step strand replacement, for which three ‘jack’ edges are added (shown in back in Figure 1j). The triangles can slide and rotate by modulating the length of the jack edges, thereby switching between the open and closed states shown in Figure 1i and j, respectively. For this purpose, we design a jack with a ‘bridge’ (a pair of crossovers of the scaffold) and two jack edges with no bridge, along with associated staple sets (see Supporting Information for design details). In the two jack edges without a bridge, the staples can be removed completely and replaced. In contrast, only a set of staples that secure the bridge (*i.e.* bridge staples) can be removed and replaced in the jack with a bridge, while other staples remain intact. In the closed state, for example, all the staples secure the scaffold segments in place such that all three jack edges are fully extended. Each staple strand for the two jacks with no bridge includes an 8-nucleotide (nt) ss-overhang which is used as a toehold for reconfiguration. In the jack edge with a bridge, only the bridge staples have the toeholds. When a set of releaser strands that are fully complementary to the staples with toeholds are added, the staples bind to the releasers and are disengaged from the scaffold, thus the staples are removed. Another set of staples can then be introduced to form a very short length for the two jack edges (*e.g.*, ~4.5 nm), while the jack with a bridge becomes undefined in length. After the second step, the origami will assume the open-state conformation. In a similar manner, the flight ring can transform from the closed to the open states.

DNA origami flight rings were assembled in a thermal cycler and visualized by AFM as shown in Figure 2 (see Supporting Information for experimental details). There are two distinct structures: equilateral triangles in the open state (Figures 2a and S4) and regular hexagons in the closed state (Figures 2b and S5). To distinguish the conformational species, the internal angle *γ* is defined, and counting in the images is presented in Figure 2c and d. The DNA nanostructures are predominantly in the designed state (~90%) with a small fraction of mis-forms and origami in the other state (~10%). The DNA flight rings in both open and closed states reasonably follow the design, although some curvature in the outer edges is observed in the open state. Each edge in the triangle subunits is on average ~51 nm in length which is designed to be approximately 49 nm or 147 base-pairs. The edge thickness also agrees well with the design. The edges in the open state have a uniform cross-section of approximately 4.5 × 4.5 nm, corresponding to four dsDNA bundles due to the overlap of the triangle subunits. The edges in the closed state have three different thicknesses: (i) With no overlap, the edges in the outer loop are two dsDNA each. (ii) Half of the inside edges have four bundles each due to two overlapped edges as shown in Figure 1j. (iii) The other three, inside edges have additional jack edges, with a total of six dsDNA each.

**Figure 2.**
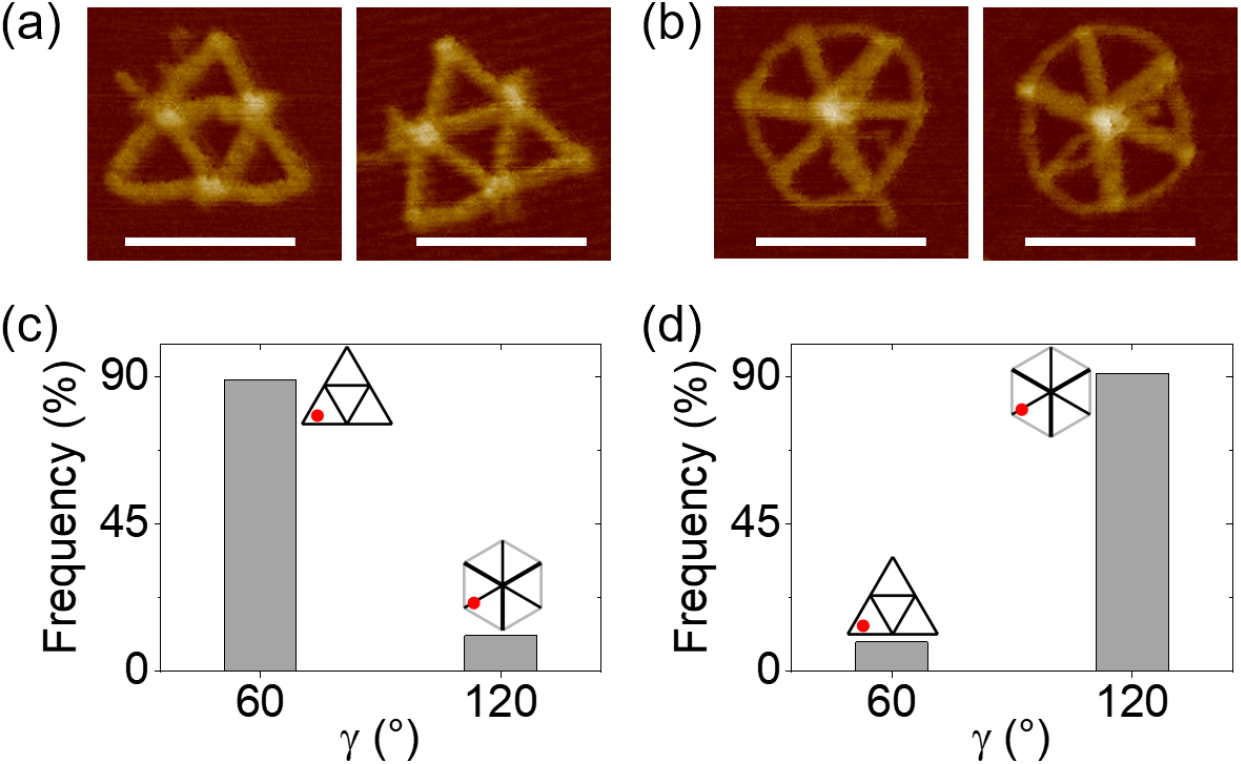
AFM images of DNA flight rings in the open (a) and the closed (b) states. Scale bar: 100 nm. While slightly curved edges are observed in the outer loop in (b), the two species are clearly distinguishable. Frequency of origami structures designed for the open state or γ = 60° (c) and the closed state or γ = 120° (d). The total numbers of DNA structures counted are 406 in (c) and 358 in (d). The inserted schematics represent the conformations with internal angle *γ* shown as red dots. The flight rings are predominantly in the designed states (~90 %).

Next, we demonstrate dynamic, deployable transformation of the DNA flight rings (Figure 3). They reconfigured from the closed to the open states as well as from the open to the closed states by changing the jack edges. As discussed above, toehold-mediated strand displacement was used to initiate the reconfiguration process (also see Supporting Information for experimental details). Figure 3a shows AFM images of the flight rings in the open state which reconfigured from the origami in the closed state (Figure 2b). The two half-length segments (~17 and 21 nm) from the jack with a bridge and the free scaffold domains (105 nt each) released from the two jacks interfere with the AFM images. Despite the blurred image, the triangle shape can be clearly identified. Similarly, the flight rings in the closed state are shown in Figure 3b which transformed from the open state (Figure 2a). The majority of the flight rings are in the designed configurations (~90%), which is comparable to the origami species in the same states before reconfiguration. The results clearly demonstrate the feasibility of deployable flight ring nanostructures from DNA.

**Figure 3.**
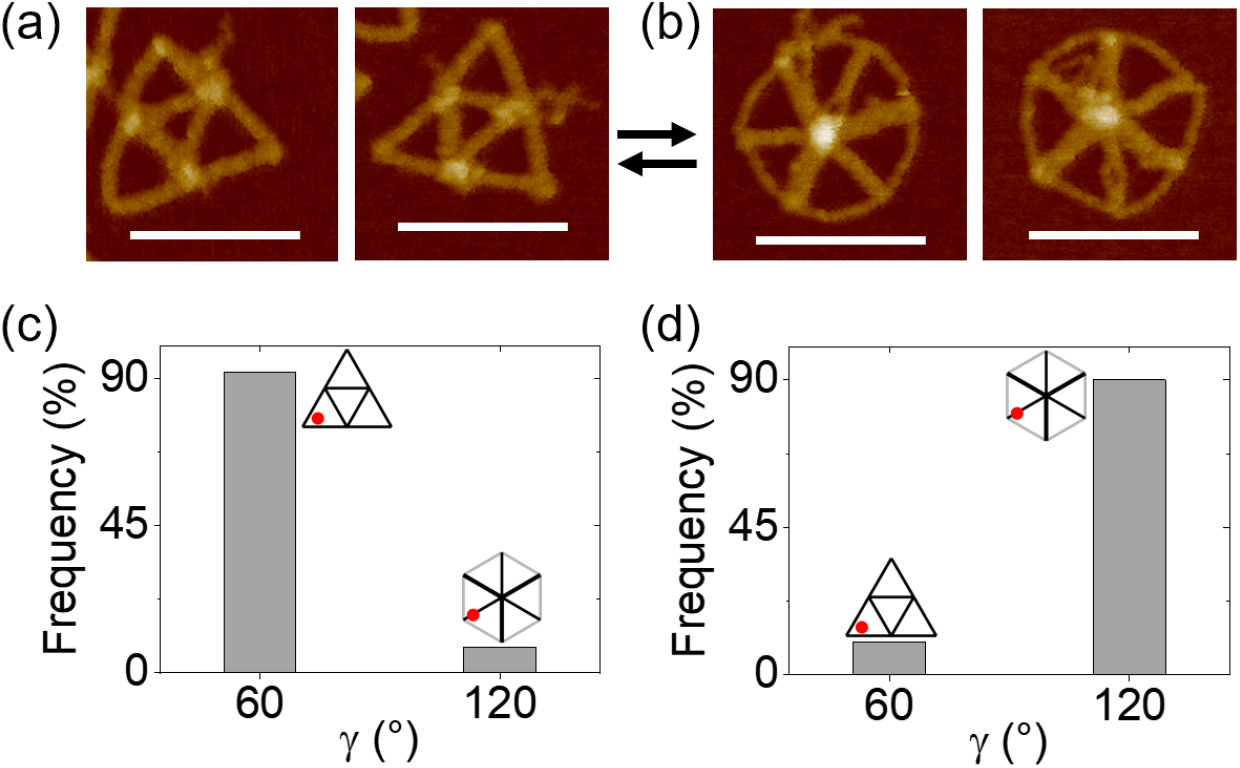
Dynamic, deployable reconfiguration of the DNA flight ring between the two states. AFM images of open (a) and closed (b) species. As the open species reconfigure from the closed state, the jack with a bridge becomes two half-length segments as the bridge staples are disengaged. The two halves and the free scaffold domains released from the two jacks with no bridge slightly blur the images. Scale bar: 100 nm. Frequency of DNA flight rings in the open (c) and closed (d) states after reconfiguration. The total numbers of DNA structures examined are 153 in (c) and 115 in (d). The inserted schematics represent the conformations with the internal angle *γ* shown as red dots. The flight rings are mostly reconfigured to designed states (~90 %), which is comparable to the origami species without reconfiguration.

Auxetic properties are key to deployable structures. To quantitatively characterize the auxetic behavior of the flight ring, Poisson’s ratio is determined. Poisson’s ratio is normally expressed as a negative ratio of strains in orthogonal directions.^27^

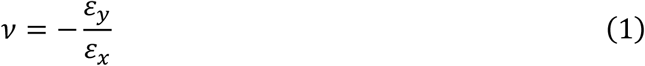

Given the shape changes between a triangle and a hexagon, however, it is not conducive to define an orthogonal direction pair in the flight ring. Instead, we develop Poisson’s ratio as a function of strains of length and area (see the derivation in Supporting Information)

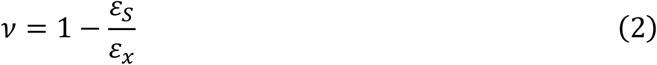

where the covered area *S* is used to define a hypothetical *y* after *x* is defined (see the inserted schematics in Figure 4).

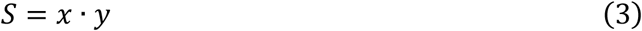

**Figure 4.**
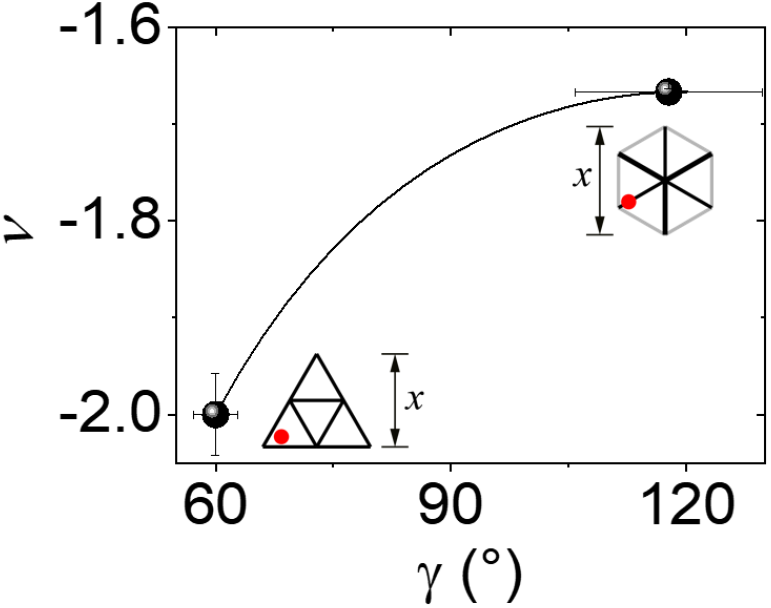
Poisson’s ratio (*ν*) versus angle (*γ*). The black line represents theoretical calculations, and the spheres are the experimental data with error bars, showing excellent agreement between the theory and experiment. The Poisson’s ratio transitions between −2.0 and −1.67, as the flight ring switches between the open (60°) and the closed (120°) states.

Considering the flight ring geometry, theoretical predictions are obtained and shown as a curve in Figure 4, which transitions between −2 (triangle) and −1.67 (hexagon).

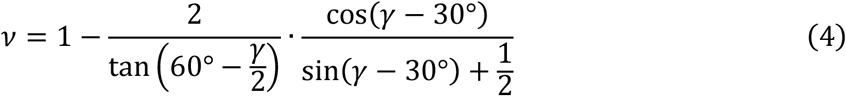

The Poisson’s ratios from the AFM images of two origami species are approximately −2.0 and −1.67 for *γ* = 60° and 120°, respectively. It is evident that the deployable transformation results in negative Poisson’s ratio as design. The flight ring is unique in that its structure is finite, while other typical auxetic geometries are periodic, extended in 2D or 3D.^27,28^

In closing, Hoberman flight rings are realized using DNA origami, which forms a left-handed trefoil knot. The deployable and auxetic nanostructures may serve as a versatile platform for topological studies and open new opportunities for bioengineering and biosensors such as high-precision drug delivery^29^ and molecular targeting and swallow^30^.

## ACKNOWLEDGEMENT

This work was financially supported by the U.S. Department of Energy. J.H.C. also acknowledges partial support from the U.S. National Science Foundation.

## REFERENCES

1 Li, R., Yao, Y.-A. & Kong, X. A Class of Reconfigurable Deployable Platonic Mechanisms. Mechanism and Machine Theory 105, 409–427 (2016).

2 Verheyen, H. F.The Complete Set of Jitterbug Transformers and the Analysis of Their Motion in Symmetry 2 Vol. 17 (ed Hargittai, I.) 203–250 (Elsevier, 1989).

3 Sun, X., Yao, Y.-A. & Li, R. Novel Method of Constructing Generalized Hoberman Sphere Mechanisms Based on Deployment Axes. Frontiers of Mechanical Engineering 15, 89–99 (2020).

4 O’Reilly, O. M. & Varadi, P. C. Hoberman’s Sphere, Euler Parameters and Lagrange’s Equations. Journal of Elasticity 56, 171–180 (1999).

5 Li, Y., Shen, Y., Cao, S., Zhang, X. & Meng, Y. Thermally Triggered Tunable Vibration Mitigation in Hoberman Spherical Lattice Metamaterials. Applied Physics Letters 114, 191904 (2019).

6 Speir, J. A., Munshi, S., Wang, G., Baker, T. S. & Johnson, J. E. Structures of the Native and Swollen Forms of Cowpea Chlorotic Mottle Virus Determined by X-Ray Crystallography and Cryo-Electron Microscopy. Structure 3, 63–78 (1995).

7 Tama, F. & Brooks III, C. L. The Mechanism and Pathway of pH Induced Swelling in Cowpea Chlorotic Mottle Virus. Journal of Molecular Biology 318, 733–747 (2002).

8 Suzuki, Y., Cardone, G., Restrepo, D., Zavattieri, P. D., Baker, T. S. & Tezcan, F. A. Self-Assembly of Coherently Dynamic, Auxetic, Two-Dimensional Protein Crystals. Nature 533, 369–373 (2016).

9 Jones, M. R., Seeman, N. C. & Mirkin, C. A. Programmable Materials and the Nature of the DNA Bond. Science 347, 1260901 (2015).

10 Wei, B., Dai, M. & Yin, P. Complex Shapes Self-Assembled from Single-Stranded DNA Tiles. Nature 485, 623–626 (2012).

11 Rothemund, P. W. Folding DNA to Create Nanoscale Shapes and Patterns. Nature 440, 297–302 (2006).

12 Castro, C. E., Kilchherr, F., Kim, D.-N., Shiao, E. L., Wauer, T., Wortmann, P., Bathe, M. & Dietz, H. A Primer to Scaffolded DNA Origami. Nature Methods 8, 221–229 (2011).

13 Marras, A. E., Zhou, L., Su, H.-J. & Castro, C. E. Programmable Motion of DNA Origami Mechanisms. Proceedings of the National Academy of Sciences 112, 713–718 (2015).

14 Dunn, K. E., Dannenberg, F., Ouldridge, T. E., Kwiatkowska, M., Turberfield, A. J. & Bath, J. Guiding the Folding Pathway of DNA Origami. Nature 525, 82–86 (2015).

15 Ke, Y., Meyer, T., Shih, W. M. & Bellot, G. Regulation at a Distance of Biomolecular Interactions Using a DNA Origami Nanoactuator. Nature Communications 7, 10935 (2016).

16 Chen, H., Zhang, H., Pan, J., Cha, T.-G., Li, S., Andréasson, J. & Choi, J. H. Dynamic and Progressive Control of DNA Origami Conformation by Modulating DNA Helicity with Chemical Adducts. ACS Nano 10, 4989–4996 (2016).

17 Tikhomirov, G., Petersen, P. & Qian, L. Fractal Assembly of Micrometre-Scale DNA Origami Arrays with Arbitrary Patterns. Nature 552, 67–71 (2017).

18 Chen, H., Li, R., Li, S., Andréasson, J. & Choi, J. H. Conformational Effects of UV Light on DNA Origami. Journal of the American Chemical Society 139, 1380–1383 (2017).

19 Kopperger, E., List, J., Madhira, S., Rothfischer, F., Lamb, D. C. & Simmel, F. C. A Self-Assembled Nanoscale Robotic Arm Controlled by Electric Fields. Science 359, 296–301 (2018).

20 You, Z. Motion Structures Extend Their Reach. Materials Today 10, 52–57 (2007).

21 Yurke, B., Turberfield, A. J., Mills, A. P., Simmel, F. C. & Neumann, J. L. A DNA-Fuelled Molecular Machine Made of DNA. Nature 406, 605–608 (2000).

22 Zhang, D. Y. & Seelig, G. Dynamic DNA Nanotechnology Using Strand-Displacement Reactions. Nature Chemistry 3, 103–113 (2011).

23 Li, Y., Chen, Y., Li, T., Cao, S. & Wang, L. Hoberman-Sphere-Inspired Lattice Metamaterials with Tunable Negative Thermal Expansion. Composite Structures 189, 586–597 (2018).

24 Sauvage, J.-P. & Dietrich-Buchecker, C. Molecular Catenanes, Rotaxanes and Knots: A Journey Through the World of Molecular Topology. (John Wiley & Sons, 2008).

25 Zhang, F., Jiang, S., Wu, S., Li, Y., Mao, C., Liu, Y. & Yan, H. Complex Wireframe DNA Origami Nanostructures with Multi-Arm Junction Vertices. Nature Nanotechnology 10, 779–784 (2015).

26 Jun, H., Wang, X., Bricker, W. P. & Bathe, M. Automated Sequence Design of 2D Wireframe DNA Origami with Honeycomb Edges. Nature Communications 10, 1–9 (2019).

27 Evans, K. E. & Alderson, A. Auxetic Materials: Functional Materials and Structures from Lateral Thinking! Advanced Materials 12, 617–628 (2000).

28 Li, R., Chen, H. & Choi, J. H. Auxetic Two-Dimensional Nanostructures from DNA. bioRxiv, doi:10.1101/2020.08.21.262139 (2020).

29 Du Plessis d’Argentré, A., Perry, S., Iwata, Y., Iwasaki, H., Iwase, E., Fabozzo, A., Will, I., Rus, D., Damian, D. D. & Miyashita, S. in 2018 IEEE International Conference on Robotics and Automation (ICRA). 1511–1518 (IEEE).

30 Liu, N., Jiang, Y., Zhou, Y., Xia, F., Guo, W. & Jiang, L. Two-Way Nanopore Sensing of Sequence-Specific Oligonucleotides and Small-Molecule Targets in Complex Matrices Using Integrated DNA Supersandwich Structures. Angewandte Chemie 125, 2061–2065 (2013).

