## Supplementary Materials for "Topological Assembly of a Deployable Hoberman Flight Ring from DNA"

**Content**

S1. Materials and Methods
S2. DNA Origami Design for the Flight Ring
S3. Poisson’s Ratio Calculation
S4. Supporting AFM Figures
S5. DNA Sequence Information
S6. References

### S1. Materials and Methods

#### Materials

All DNA oligomers (including staples and releasers) were obtained from Integrated DNA Technologies (sequence information presented in S5). The M13mp18 scaffold was supplied by Bayou Biolabs. All DNA strands were used directly from the plates or tubes without further purification. All other chemicals were purchased from Sigma Aldrich.

#### DNA origami synthesis

Flight rings were synthesized by mixing 5 nM scaffold strands with 4× DNA staple strands in 1× TAE buffer (an aqueous solution of 40 mM trisaminomethane, 1 mM ethylenediaminetetraacetic acid (EDTA) disodium salt, and 20 mM acetic acid at pH ~8) that also contains 6 mM magnesium acetate (termed TAEM6 buffer). The mixture was then thermally annealed from 95 to 4 °C in a process described as follows. First, the mixture was annealed from 95 °C to 65 °C at -1 °C per 2 min; then it went from 65 °C to 60 °C at -1 °C per 25 min; from 60 °C to 50 °C at -1 °C per 60 min; from 50 °C to 45 °C at -1 °C per 25 min; from 45 °C to 25 °C at -1 °C per 2 min; at the end, it was then held at 4 °C. The annealing was performed with a BIO-RAD S1000 Thermal Cycler. No further purification was performed before AFM imaging.

#### DNA origami reconfiguration

There are two major steps in the reconfiguration: strand displacement and reannealing. The flight rings from synthesis were mixed with 10× releaser strands. The mixture was then incubated at 55 °C or 60 °C for 1 hour for toehold-mediated strand displacement. After that, purifications were performed 3 times by using the centrifugal filter (100 kDa) from Amicon. Then, 20× new DNA staples for the released domain were mixed with the purified structures and incubated at 45 °C for 1 hour. The mixture was then annealed from 45 °C to 20 °C at a rate of -1 °C per 1 min. After the incubation, the mixture was held at 4 °C. Both the incubation and reannealing were performed with a BIO-RAD S1000 Thermal Cycler. No further purification was performed before AFM imaging.

#### AFM imaging

For deposition, the target sample was first diluted to 0.5 to 1.0 nM with TAEM6 buffer. An aliquot of 10 µL diluted sample was pipetted onto mica surface and incubated for 5 min at room temperature. Then, 20 µL TAEM6 buffer that also contains 2 mM nickel chloride was added onto the mica to increase the adhesion of DNA to mica. After 2 min of second incubation at room temperature, the mica was blown dry with compressed air, rinsed with 80 µL deionized (DI) water for about 3 sec, and then blown dry again with compressed air.

AFM imaging was performed in air in the Peak-Force tapping mode with a Bruker Dimension Icon AFM and SCANASYST-AIR probes. For each mica surface, at least three different places were imaged. In each given place, the scanning scales were 5 × 5 µm, 2 × 2 µm, and 500 × 500 nm.

**S2. Design of Flight Ring**

**Flat and topological designs**

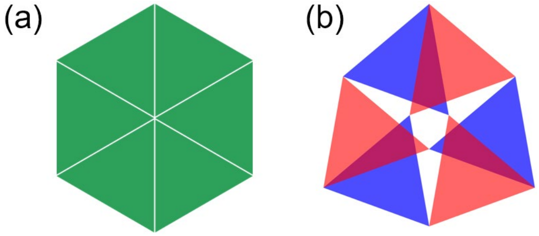


**Figure S1.** Schematics of flat (a) and topological (b) designs. (a) A single-layer hexagon is formed by six green triangles. (b) A flight ring is constructed with six triangles. Red triangles are on the top of blue triangles. The vertex of each red triangle is connected to that of a blue triangle on the opposite side, thus forming a topological design.

Figure S1 compares a topological design and a typical flat design. For example, a flat hexagon is formed by six triangles on a single plane, and its design process is straightforward. In contrast, a flight ring has a topological design with six triangles such that half of the triangles (shown in red) are on top of the other three (blue). The inside vertex of a triangle is connected to that of the triangle in different color (*i.e.*, different layer) on the opposite side. In this topological design (Figure S1b), the blue and red triangles in two layers can slide against each other, thus transforming the structure between two distinct (open and closed) states.

**Handedness of trefoil knots**

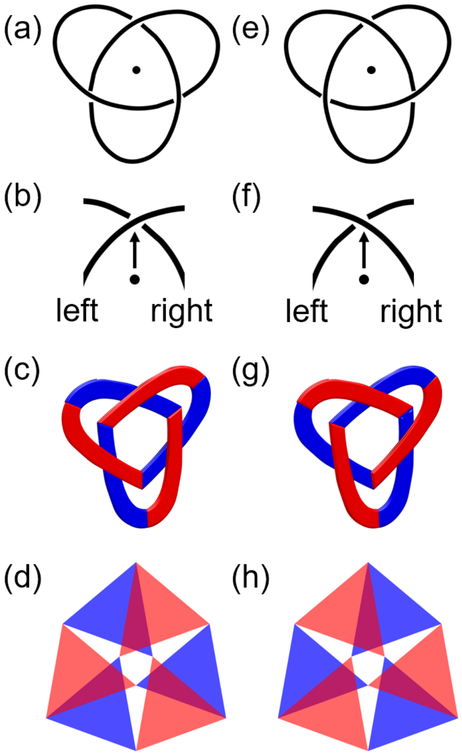


**Figure S2.** Schematics of trefoil knots. (a) – (c) Topological demonstration of a left-handed trefoil knot with the center marked as a black dot in (a) – (b). The arrow in (b) indicates the direction pointing from the center to the crossing point. (d) A flight ring may be simplified into a left-handed trefoil knot. (e) – (g) Topological demonstration of a right-handed trefoil knot with the center marked as a black dot in (e) – (f). The arrow in (f) indicates the direction pointing from the center to the crossing point. (h) A flight ring can be simplified to a right-handed knot. Red represents the top layer, while blue denotes the bottom layer.

Trefoil knots have left-handed (Figure S2a – d) or right-handed species (Figure S2e – h). The difference is manifested by the order of the routes. In the left-handed knot, for example, the left-side route always goes over the right-side route at crossing points with respect to the direction from the center to the crossing point (Figure 2a – c). In the right-handed knot, the arrangement is the opposite (Figure 2e – g).

**Routing of the scaffold strand and arrangement of staples**

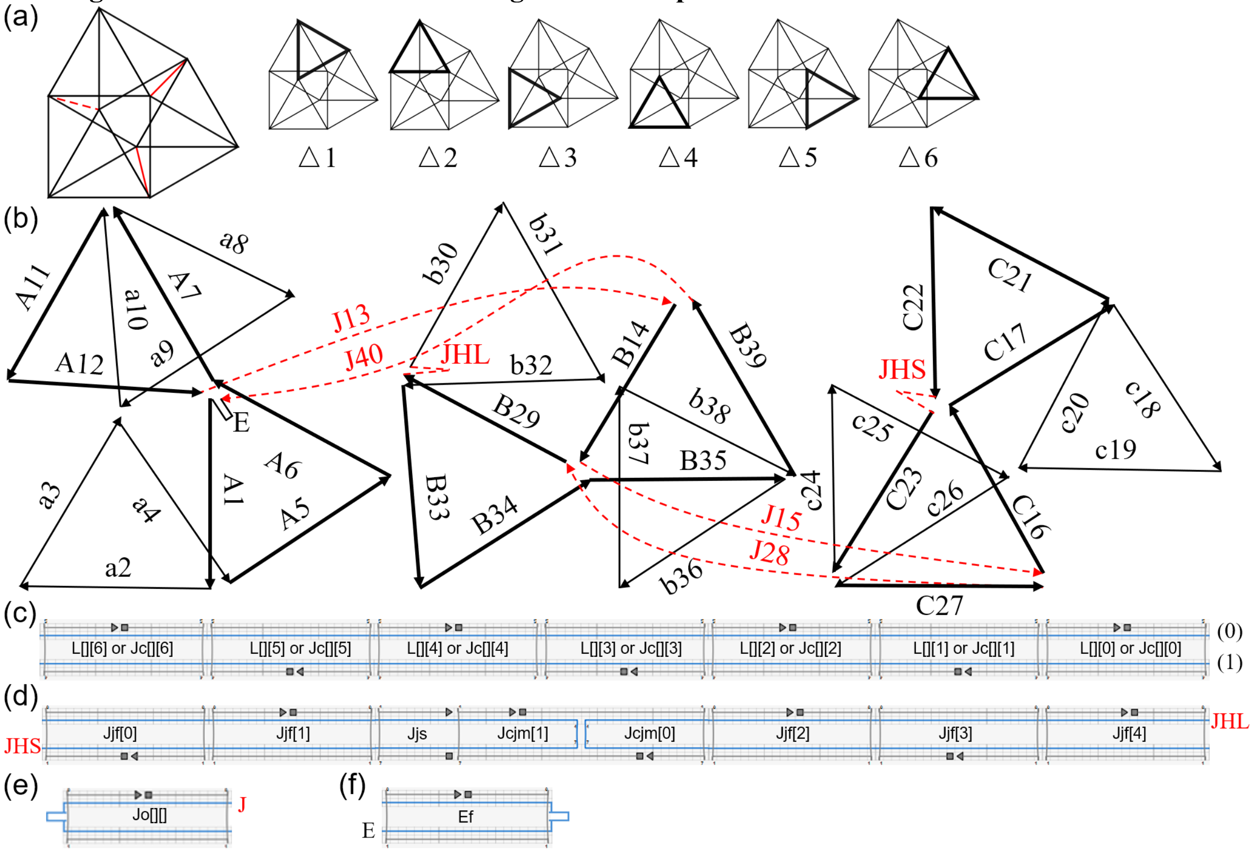


**Figure S3.** Schematics of scaffold routing and staple arrangement in the flight ring design. (a) Overall design with six triangles (△1 to △6) marked as bold lines. Black lines represent regular edges (total 18), while jack edges are shown in red (total 3). The jack on the left, shown as a red dashed line, is designed with a pair of crossovers of the scaffold (*i.e.* bridge). The other two jack edges (on the right and on the bottom) do not have a bridge, which are shown as solid red lines. (b) Routing of the scaffold. The arrows denote the direction of the scaffold from 5' to 3' end. Due to overlapping, the routing is demonstrated by three parts, A, B and C, for clarification. Lowercase letters represent the three contiguous scaffold segments (shown as thin lines) for a triangle; uppercases indicate another three scaffold segments for a triangle (bold lines) or jacks that connect triangles (red dashed lines). Each triangle is routed twice, one with thin lines and the other with bold lines, in antiparallel. Numbers denote the order of the scaffold segments, starting from A1 and ending at J40. In addition, the 'E' between A1 and J40 is designed to stabilize the extra scaffold segment. A1 through J40 are grouped into L[0] to L[17] (regular edges), J[0] and J[1] (jack edges) with detailed information presented in Table S1. (c) – (f) The detailed edge designs in Cadnano2.[1] Blue lines represent the scaffold segments, and staples are shown in gray. Black squares and triangles mark the 5' and 3' ends of the staples, respectively. The numbers to be filled in the square brackets are the number of the corresponding edge. For example, the left end staple of edge L[] is L[][6]. For the edge L[0], this staple is called L[0][6]. (c) All regular edges (L[0] to L[17]), and two jack edges with no bridge (J[0] and J[1], shown as solid red lines on the right and bottom in (a), respectively) when fully extended. In each edge, the two scaffold segments are recognized as (0) and (1) as listed in Table S1. These two, upper and lower scaffold segments, are 147 nt each. The staples, L[][0] to L[][6] (or Jc[][0] to Jc[][6]), are 42 nt each. The staple name starts with 'L' for a regular edge, while it starts with 'Jc' for a jack edge. (d) The jack edge with a bridge when fully extended. This jack, which consists of JHL and JHS, is named JB and shown as the red dashed line on the left in (a). The left scaffold segment is JHS (136 nt), and the right is JHL (158 nt). Both are labelled at the 5' ends. The staples, Jjs, Jcjm[0], Jcjm[1], and Jjf[0] to Jjf[4], have different lengths, varying from 22 to 42 nt. Jcjm[0] and Jcjm[1] are marked at the 5' ends. (e) Two jack edges, J[0] and J[1], when fully shrunk. The free, unpaired scaffold loop, marked as a bump in blue, on the left is 105 nt. The scaffold segment (147 nt) in the design is applicable for both segments of each jack. For example, in J[1], segment (0) is J15; in this case, the staple name is Jo[1][0] which is 42 nt. (f) The stabilizer for the extra scaffold segment. The scaffold segment, named E, is 768 nt. The free loop, marked as a blue bump on the right, is 726 nt. The staple Ef is 42 nt.

Since the scaffold strand goes through two layers that can slide against each other in the flight ring structure, it is important to have as few interferences between triangles as possible. The pair of crossovers of the scaffold (*i.e.* bridge) may create a possible issue. For example, if two of the three edges of a triangle are formed and there is a bridge on the third, unformed edge, the triangle may lock with another triangle due to the bridge. Thus, a bridge is designed in one of the jack edges named JB (between the scaffold segments JHS and JHL as shown in Figure S3d) to avoid a locking of triangle sliding. The names and segments of edges without a bridge (all regular edges, L[0] to L[17], and two jack edges, J[0] and J[1]) are specified in Table S1.

**Table S1.** Names and segments of edges without a bridge.

| name | segment (0) | segment (1) |
|---|---|---|
| L[0] | A1 | b37 |
| L[17] | A5 | b36 |
| L[9] | A6 | b38 |
| L[10] | A7 | b31 |
| L[2] | A11 | b30 |
| L[1] | A12 | b32 |
| J[0] | J13 | J40 |
| L[7] | B14 | c20 |
| J[1] | J15 | J28 |
| L[13] | C16 | a4 |
| L[12] | C17 | a9 |
| L[11] | C21 | a8 |
| L[3] | C22 | a10 |
| L[4] | C23 | a3 |
| L[14] | C27 | a2 |
| L[6] | B29 | c25 |
| L[5] | B33 | c24 |
| L[15] | B34 | c26 |
| L[16] | B35 | c19 |
| L[8] | B39 | c18 |

The designed length of regular edges (*l*) are calculated to be approximately 49 nm based on 0.332 nm/bp along the axis of regular, right-handed B-DNA. The length of the jack edges is modulated. When fully extended, the length of all three jacks is ~49 nm (the same as the regular edges) and the flight ring is in the closed state. When fully shrunk, the length of a jack edge is theoretically 0 nm and the origami structure is in the open state. In our experiment, the length of two jacks J[0] and J[1] is approximately 4.5 nm (*i.e.*, the thickness of 2-dsDNA helices) in the closed state (when fully shrunk). On the other hand, the length of the jack JB becomes undefined as the staples for the bridge (*i.e.*, bridge staples, Jcjm[0] and Jcjm[1]) are not provided. JHS and JHL, the two parts of disengaged JB, are ~17 and 21 nm, respectively. To initiate structural transformation, toeholds are placed at the 3' end of 20 jack staples (Jc[0][0] through Jc[1][6], Jcjm[0], Jcjm[1], and Jo[0][0] through Jo[1][1]) (see S5 for details).

The joints linking the edges together are also important, which must maintain a balance between flexibility and structural integrity. To simplify the design, all the joints are made of single-stranded scaffold segments and the lengths are adjusted based on whether the angle of the edges that connect to the joint is changeable. For example, the angle between a2 and a3 is unchangeable as they form the triangle 4 or △4; in this joint a short ssDNA is designed such as 3 nt. In contrast, the angle between a4 and A5 may change, since they connect △4 and △5, which can slide against each other. Here, a longer ssDNA segment is given (*e.g.*, 10 nt). All the joints are referred by the ending scaffold segments (from 5' to 3' end) – for example, the joint from a2 to a3 is noted as a3 (see the list below). The joints that connect to the stabilizer (E) are exceptions and are different from others, which as shown below as A1 and E.

- 3 nt: a3, a4, A6, a9, a10, A12, c19, c20, C22, c25, c26, b31, b32, B34, b37, b38
- 4 nt: A1
- 5 nt: E
- 10 nt: a2, A5, A7, a8, A11, J13, B14, J15, C16, C17, c18, C21, JHS, C23, c24, C27, J28, B29, JHL, b30, B33, B35, b36, B39, J40

**Validation of topological design**

From the design details, the knot type may be analyzed. For any two triangles, if they share a vertex and overlap with each other, they must be in different layers. For example, △1 consists of a8 to a10, C17, C21 and C22, while △2 consists of A7, A11, A12 and b30 to b32. The visual inspection of A7 to A12 indicates that △1 and △2 share the point where a8 meets a10 and where A7 meets A11. In addition, △1 and △2 overlap with each other. Thus, △1 and △2 are in different layers. A similar analysis may be performed to conclude that △5 and △6, as well as △3 and △4 are two groups that are in different layers. If we use "=" to represent being in the same layer and "≠" for different layers.

$$\triangle 1 \neq \triangle 2 \quad (2.1)$$

$$\triangle 5 \neq \triangle 6 \quad (2.2)$$

$$\triangle 3 \neq \triangle 4 \quad (2.3)$$

Another analysis may be used for the triangles that do not overlap with each other. If any two triangles share the same vertex and there is a cross-point of the edges, the cross-point will prevent the triangles from sharing the same layer. A good example is △4 and △5 which share the vertex at the place where a2 meets a4 and where A1 meets A5. There is a cross-point between A1 and a4. Therefore, △4 and △5 are in different layers. Similarly, △1 and △6, as well as △2 and △3 are two groups in different layers.

$$\triangle 1 \neq \triangle 6 \quad (2.4)$$

$$\triangle 2 \neq \triangle 3 \quad (2.5)$$

$$\triangle 4 \neq \triangle 5 \quad (2.6)$$

Given that there are only two layers,

$$\triangle 1 = \triangle 3 = \triangle 5 \neq \triangle 2 = \triangle 4 = \triangle 6 \quad (2.7)$$

Therefore, this design will have either left-handed trefoil knot (Figure S2c) or right-handed trefoil knot (Figure S2e), depending on whether △1 is on the top or bottom.

**S3. Poisson's Ratio Calculation**

From the geometry in insets of Figure 4, length $x$ and area $S$ are expressed.

$$x = 2l\cos\left(\frac{\pi}{3} - \frac{\gamma}{2}\right) \tag{3.1}$$

$$S = \sqrt{3}l^2\left[\sin\left(\gamma - \frac{\pi}{6}\right) + \frac{1}{2}\right] \tag{3.2}$$

Displacements and strains are then derived.

$$\mathrm{d}x = l\sin\left(\frac{\pi}{3} - \frac{\gamma}{2}\right)\mathrm{d}\gamma \tag{3.3}$$

$$\varepsilon_x = \frac{\mathrm{d}x}{x} = \frac{\tan\left(\frac{\pi}{3} - \frac{\gamma}{2}\right)}{2}\mathrm{d}\gamma \tag{3.4}$$

$$\mathrm{d}S = \sqrt{3}l^2\cos\left(\gamma - \frac{\pi}{6}\right)\mathrm{d}\gamma \tag{3.5}$$

$$\varepsilon_S = \frac{\mathrm{d}S}{S} = \frac{\cos\left(\gamma - \frac{\pi}{6}\right)}{\sin\left(\gamma - \frac{\pi}{6}\right) + \frac{1}{2}}\mathrm{d}\gamma \tag{3.6}$$

Poisson's ratio may be expressed as a negative ratio of strains in transverse directions.

$$\nu = -\frac{\varepsilon_y}{\varepsilon_x} \tag{3.7}$$

where

$$\varepsilon_y = \frac{\Delta y}{y} \tag{3.8}$$

However, it is difficult to define $x$ and $y$ simultaneously in the flight ring. Thus, a hypothetical $y$ may be expressed by $x$ and $S$.

$$S = x \cdot y \tag{3.9}$$

$$S + \Delta S = (x + \Delta x) \cdot (y + \Delta y) \tag{3.10}$$

Then,

$$S(1 + \varepsilon_S) = x(1 + \varepsilon_x) \cdot y(1 + \varepsilon_y) \tag{3.11}$$

$$\varepsilon_S = \varepsilon_x + \varepsilon_y + \varepsilon_x \cdot \varepsilon_y \tag{3.12}$$

With the assumption of small $\varepsilon_x$ and $\varepsilon_y$,

$$\varepsilon_S \cong \varepsilon_x + \varepsilon_y = (1 - \nu) \cdot \varepsilon_x \tag{3.13}$$

Thus,

$$\nu = 1 - \frac{\varepsilon_S}{\varepsilon_x} \tag{3.14}$$

Combining equation (3.4) and (3.6),

$$\nu = 1 - \frac{2}{\tan\left(\frac{\pi}{3} - \frac{\gamma}{2}\right)} \cdot \frac{\cos\left(\gamma - \frac{\pi}{6}\right)}{\sin\left(\gamma - \frac{\pi}{6}\right) + \frac{1}{2}} \tag{3.15}$$

### S4. Supporting AFM Figures

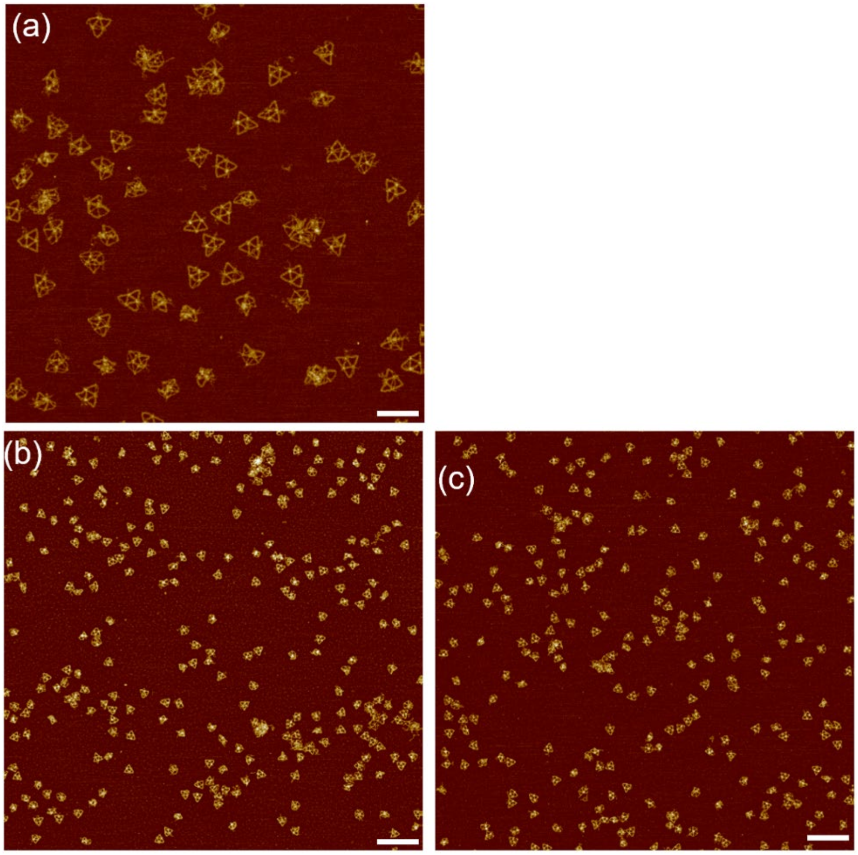


**Figure S4.** AFM images of DNA origami flight rings designed for the open state. The total structures counted are 406 with 361 in open and 45 in closed states. Scale bar: (a) 100 nm; (b) – (c) 250 nm.

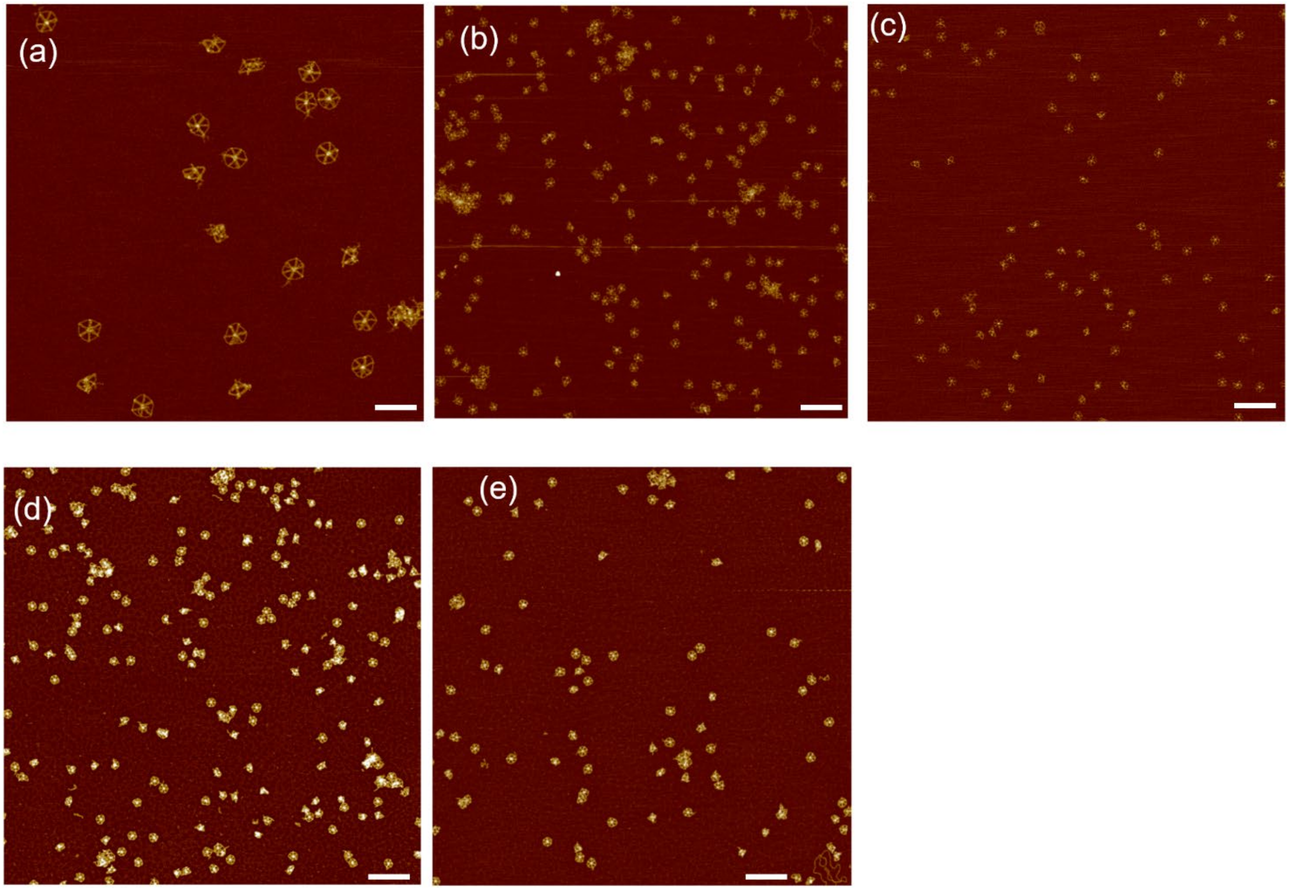


**Figure S5.** AFM images of DNA origami flight rings designed for the closed state. The total structures counted are 358 with 32 in open and 326 in closed states. Scale bar: (a) 100 nm; (b) – (e) 250 nm.

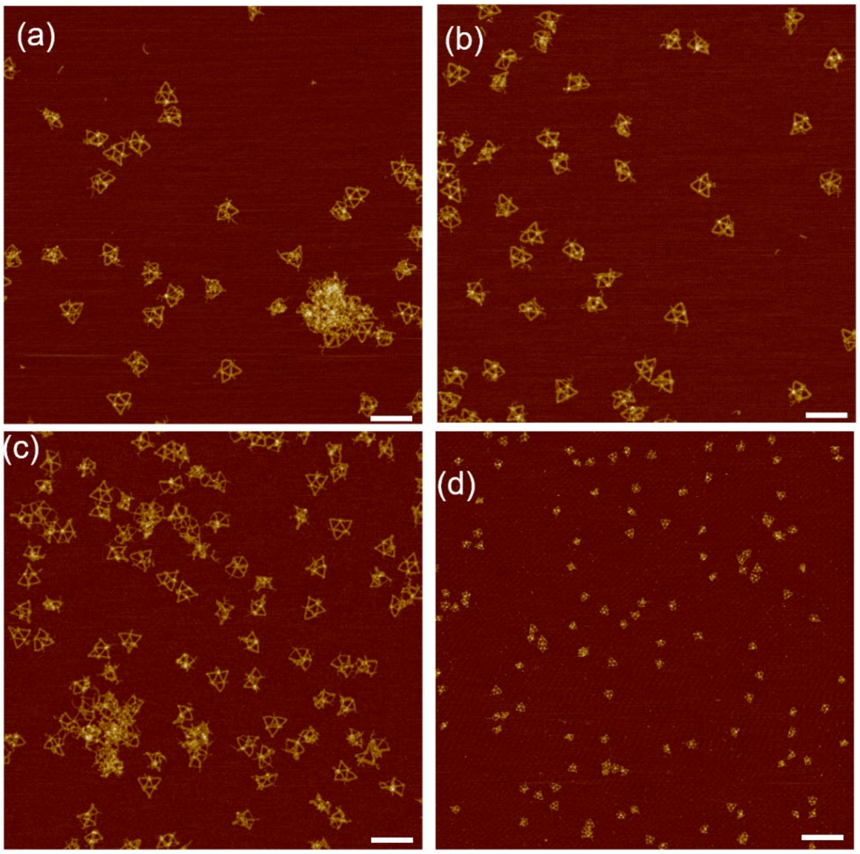


**Figure S6.** AFM images of DNA origami flight rings after reconfiguration from the closed state. The total structures counted are 153 with 141 in open and 12 in closed states. Scale bar: (a) – (c) 100 nm; (d) 250 nm.

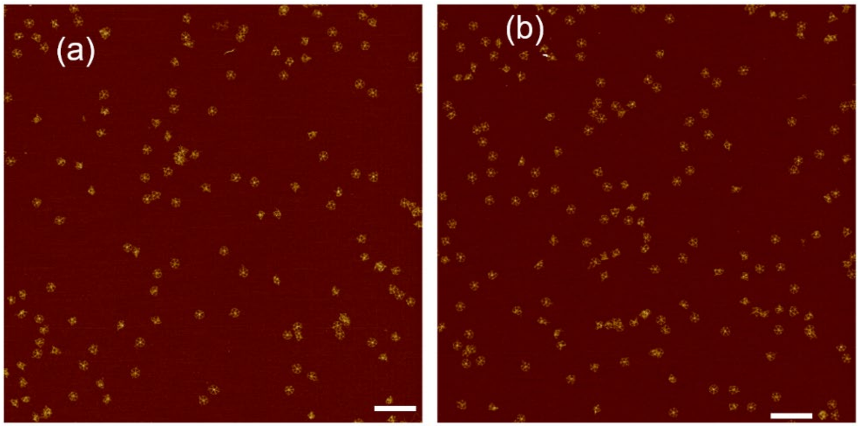


**Figure S7.** AFM images of DNA origami flight rings after reconfiguration from the open state. The total structures counted are 115 with 20 in open and 95 in closed states. Scale bar: 250 nm.

**S5. DNA Sequence Information**

All the staple sequences are listed below. The releaser strands for reconfiguration are also presented.

| Name | Sequence (5' to 3') of Staple Strand |
|---|---|
| L[0][0] | GTAGTAGCATTTTGGGCGCCAGGGTGGTTTTTCTACTAATA |
| L[0][1] | GTGAGACGGGCAGCTGAAAAGGTGGCATCAATCTTTTCACCA |
| L[0][2] | TTTGGGGCGCGAACAGCTGATTGCCCTTCACCTATATTTTCA |
| L[0][3] | GAGAGAGTTGCATGGTCAATAACCTGTTTAGCGCCTGGCCCT |
| L[0][4] | ACATTTCGCAAAGCAAGCGGTCCACGCTGGTTACCATTAGAT |
| L[0][5] | GGCGAAAATCCCGAACGAGTAGATTTAGTTTGTGCCCCAGCA |
| L[0][6] | TCCCAATTCTGTGTTTGATGGTGGTTCCGAAAAACAGTTGAT |
| L[1][0] | TTCCAGACGTTTTGCTGGTAATATCCAGAACATTTGTCGTCT |
| L[1][1] | CAGCCATTGCAGTTAGCGTAACGATCTAAAGTATATTACCGC |
| L[1][2] | CAGCCCTCATAACAGGAAAAACGCTCATGGAAATTCCACAGA |
| L[1][3] | TTTGACGCTCAACAAACTACAACGCCTGTAGCATACCTACAT |
| L[1][4] | TCGTCACCAGTATCGTCTGAAATGGATTATTTACACTGAGTT |
| L[1][5] | ATTCACCAGTCAATAGGAACCCATGTACCGTAACATTGGCAG |
| L[1][6] | ATAGCAAGCCCACACGACCAGTAATAAAAGGGATTTTCAGGG |
| L[2][0] | GGAGCCTTTAAGATAAAACAGAGGTGAGGCGGGGCTCCAAAA |
| L[2][1] | CACCGCCTGCAGTTGAAAATCTCCAAAAAAAAATCAGTATTAA |
| L[2][2] | AATTTTTTCACACAGTGCCACGCTGAGAGCCATGCGAATAAT |
| L[2][3] | AAAAATCTAAAGAAAGGAACAACTAAAGGAATGCAGCAAATG |
| L[2][4] | GAGTGAGAATAGCATCACCTTGCTGAACCTCAAGTTTCAGCG |
| L[2][5] | CCTCAATCAATTTTTGCTAAACAACTTTCAACAATATCAAAC |
| L[2][6] | TCTGTATGGGAATCTGGTCAGTTGGCAAATCAAAATGAATTT |
| L[3][0] | AACGAGCGTCTTTATCAGCTTGCTTTCGAGGTTACCAACGCT |
| L[3][1] | AACAGCTTGATCTACAATTTTATCCTGAATCTGAATTTCTTA |
| L[3][2] | TTTGCACCCAGACCGATAGTTGCGCCGACAATTAGTTGCTAT |
| L[3][3] | ATCGCCCACGCTTTGAAGCCTTAAATCAAGATGACAACAACC |
| L[3][4] | TTGCGGGAGGTATAACCGATATATTCGGTCGCACCTCCCGAC |
| L[3][5] | AGGGAGTTAAAGAACGCGAGGCGTTTTAGCGATGAGGCTTGC |
| L[3][6] | CGGTATTCTAAGGCCGCTTTTGCGGGATCGTCAAGGCTTATC |
| L[4][0] | TCCTAATTTACATGACCATAAATCAAAAATCATAATATCCCA |
| L[4][1] | CTGACTATTATATAAGTCCTGAACAAGAAAAAGGTCTTTACC |
| L[4][2] | ATCAACAATAGAGTCAGAAGCAAAGCGGATTGGCGCCTGTTT |
| L[4][3] | ATTAAGAGGAACATGTTCAGCTAATGCAGAACCATCAAAAAG |
| L[4][4] | ACAATAAACAAGCCCGAAAGACTTCAAATATCCCAGACGACG |
| L[4][5] | TCGAGCTTCAACAAAAGGTAAAGTAATTCTGTGCGTTTTAAT |
| L[4][6] | TAAAGTACCGAAGCGAACCAGACCGGAAGCAATAAGAGAATA |

| | |
|---|---|
| L[5][0] | GAAGAACTCAAGAATCATAATTACTAGAAAAATGCCTGAGTA |
| L[5][1] | TATCATATGCGTTGATTAGTAATAACATCACTGCCTGTTTAG |
| L[5][2] | GCAATACTTCTTTATACAAATTCTTACCAGTAAACCGTTGTA |
| L[5][3] | GCTCAACAGTAAGTCTGTCCATCACGCAAATTTAAAGCCAAC |
| L[5][4] | CCGAGTAAAAGGGGCTTAATTGAGAATCGCCAAGTGAGGCCA |
| L[5][5] | CGCCAACATGTCTGAGAAGTGTTTTTATAATCTATTTAACAA |
| L[5][6] | ACGCCAGAATCAATTTAGGCAGAGGCATTTTCCAGGAACGGT |
| L[6][0] | AGAACCTACCAGTAAATGCTGATGCAAATCCATGGAAGGGTT |
| L[6][1] | AAAGAACGCGATTGGATTATACTTCTGAATAAATCGCAAGAC |
| L[6][2] | ATCCTGATTGTGAAAACTTTTTCAAATATATTCATCAATATA |
| L[6][3] | TCATCTTCTGAGATTATCAGATGATGGCAATTTTAGTTAATT |
| L[6][4] | ATCATATTCCTCCTAAATTTAATGGTTTGAAACGGAATTATC |
| L[6][5] | GTGATAAATAACAAAGAAACCACCAGAAGGAGTACCGACCGT |
| L[6][6] | ATTTTGCGGAAGGCGTTAAATAAGAATAAACATAACATTATC |
| L[7][0] | CCTCAAGAGAAGAACACCCTGAACAAAGTCAGGGCTGAGACT |
| L[7][1] | AGCGCTAATATTGAAACATGAAAGTATTAAGAAGGGTAATTG |
| L[7][2] | ACCTATTATTCCAGAGAGATAACCCACAAGAACTATTTCGGA |
| L[7][3] | CCCAATAATAATAAACAGTTAATGCCCCCTGCTTGAGTTAAG |
| L[7][4] | CAGTGCCCGTAGAGCAAGAAACAATGAAATAGCCTTGAGTAA |
| L[7][5] | CTTACCGAAGCTAAGTTTTAACGGGGTCAGTGCAATAGCTAT |
| L[7][6] | TGTACTGGTAACCTTTTTAAGAAAAGTAAGCAGATACAGGAG |
| L[8][0] | TTCCACACAACAAAGACACCACGGAATAAGTTCCGCTCACAA |
| L[8][1] | CAATCAATAGATTTCCTGTGTGAAATTGTTATTATTTTGTCA |
| L[8][2] | GGTCATAGCTGAAATTCATATGGTTTACCAGCTCGTAATCAT |
| L[8][3] | AAAGGGCGACACCCCGGGTACCGAGCTCGAATGCCAAAGACA |
| L[8][4] | CTCTAGAGGATTTCAACCGATTGAGGGAGGGATGCAGGTCGA |
| L[8][5] | TGACGGAAATTCCAGTGCCAAGCTTGCATGCCAGGTAAATAT |
| L[8][6] | TAAAACGACGGATTCATTAAAGGTGAATTATCCACGACGTTG |
| L[9][0] | CTACGTTAATAGAAGCATAAAGTGTAAAGCCTGAAGAAAAAT |
| L[9][1] | ATGAGTGAGCTTTATACCAGTCAGGACGTTGGGGGGTGCCTA |
| L[9][2] | GAACTGGCTCAAACTCACATTAATTGCGTTGCGCGATTTTAA |
| L[9][3] | CGCTTTCCAGTAATCATTGTGAATTACCTTATGCTCACTGCC |
| L[9][4] | ATTTCAACTTTCGGGAAACCTGTCGTGCCAGCAGATGGTTTA |
| L[9][5] | AATCGGCCAACACGAGTAGTAAATTGGGCTTGTGCATTAATG |
| L[9][6] | GAAACACCAGAGCGCGGGGAGAGGCGGTTTGCGCCCTGACGA |
| L[10][0] | CTCATTCAGTGTTCTGGCCAACAGAGATAGAAAACAAAGCTG |
| L[10][1] | CTGAAAGCGTAATTCATTACCCAAATCAACGTCCCTTCTGAC |
| L[10][2] | AAGAACCGGATAGAATACGTGGCACAGACAATTAATCTTGAC |

| | |
|---|---|
| L[10][3] | GGCTATTAGTCCTGGCTGACCTTCATCAAGAGATTTTTGAAT |
| L[10][4] | AGGCGCATAGGTTTAATGCGCGAACTGATAGCTGTACAGACC |
| L[10][5] | CGCCATTAAAATGAAAGAGGACAGATGAACGGCCTAAAACAT |
| L[10][6] | CTGACCAACTTATACCGAACGAACCACCAGCAGGGAACCGAA |
| L[11][0] | CATTAGACGGGGAATACACTAAAACACTCATCCAGGGAAGCG |
| L[11][1] | AGCGATTATACACAGAGAGAATAACATAAAAATTTGACCCCC |
| L[11][2] | TAGCAGCCTTTCAAGCGCGAAACAAAGTACAAAAAATGAAAA |
| L[11][3] | TATCATCGCCTACGATTTTTTGTTTAACGTCACGGAGATTTG |
| L[11][4] | CCAAATAAGAAGATAAATTGTGTCGAAATCCGTTATCCCAAT |
| L[11][5] | CATGTTACTTACAAAATAAACAGCCATATTATCGACCTGCTC |
| L[11][6] | TTTGCCAGTTAGCCGGAACGAGGCGCAGACGGCAGAGCCTAA |
| L[12][0] | CGTCAGACTGTCTCAGCAGCGAAAGACAGCATTTGCCTTTAG |
| L[12][1] | GTAGCAACGGCTCAGTAGCGACAGAATCAAGTCGGAACGAGG |
| L[12][2] | CAGCACCGTAATACAGAGGCTTTGAGGACTAACCATCGATAG |
| L[12][3] | ATGAGGAAGTTCCGGAAACGTCACCAATGAAAAGACTTTTTC |
| L[12][4] | CATTAGCAAGGTCCATTAAACGGGTAAAATACGCACCATTAC |
| L[12][5] | TACGAAGGCACGAGCCAGCAAAATCACCAGTAGTAATGCCAC |
| L[12][6] | TTTGGGAATTACAACCTAAAACGAAAGAGGCAACTTGAGCCA |
| L[13][0] | ACCCTCAGAGCAACCAAAATAGCGAGAGGCTTCAGAGCCACC |
| L[13][1] | AGTTTTGCCAGCACCCTCAGAACCGCCACCCTTTGCAAAAGA |
| L[13][2] | CTCAGAGCCGCAGGGGGTAATAGTAAAATGTTAACCGCCTCC |
| L[13][3] | AGCGTCCAATACGGAACCAGAGCCACCACCGGTAGACTGGAT |
| L[13][4] | ATCAAAATCACCTGCGGAATCGTCATAAATATTCTTTTCATA |
| L[13][5] | CCCCTCAAATGCCCCCTTATTAGCGTTTGCCATCATTGAATC |
| L[13][6] | TTCGGTCATAGCTTTAAACAGTTCAGAAAACGCATCGGCATT |
| L[14][0] | CCTTGCTTCTGCCAACAGGTCAGGATTAGAGAGAGTGAATAA |
| L[14][1] | TTGCTCCTTTTAGTACATAAATCAATATATGTGTACCTTTAA |
| L[14][2] | TTAATGGAAACGATAAGAGGTCATTTTTGCGGATTACCTTTT |
| L[14][3] | GCTTAATTGCTACATTTAACAATTTCATTTGAATGGCTTAGA |
| L[14][4] | CAAAATTAATTGAATATAATGCTGTAGCTCAAATCAAGAAAA |
| L[14][5] | ATATGCAACTAAAAGAAGATGATGAAACAAACCATGTTTTAA |
| L[14][6] | TTACCTGAGCAAAGTACGGTGTCTGGAAGTTTTTCATTTCAA |
| L[15][0] | AAAGGGATTTTCTATTAATTAATTTTCCCTTAGAGGCCGATT |
| L[15][1] | AAACATAGCGAATCAGAGCGGGAGCTAAACAGGAATCCTTGA |
| L[15][2] | TCCTCGTTAGATAGCTTAGATTAAGACGCTGATAACGTGCTT |
| L[15][3] | TAGTGAATTTAGGTTGCTTTGACGAGCACGTAGAAGAGTCAA |
| L[15][4] | GCGCGTACTATTCAAAATCATAGGTCTGAGAGCCGCTACAGG |
| L[15][5] | TAACCTCCGGCACACCCGCCGCGCTTAATGCGACTACCTTTT |

| | |
|---|---|
| L[15][6] | GCGTAACCACCTTAGGTTGGGTTATATAACTAGTCACGCTGC |
| L[16][0] | AGGGCGCTGGCAGCCGAACAAAGTTACCAGAAAGCGGGCGCT |
| L[16][1] | GAAACGCAATAGGAAGGGAAGAAAGCGAAAGGGGAAACCGAG |
| L[16][2] | GTGGCGAGAAAATAACGGAATACCCAAAAGAAGCCGGCGAAC |
| L[16][3] | TAAGACTCCTTATTTAGAGCTTGACGGGGAAACTGGCATGAT |
| L[16][4] | GGGAGCCCCCGATTACGCAGTATGTTAGCAAAGAACCCTAAA |
| L[16][5] | ACATACATAAAGTGCCGTAAAGCACTAAATCGCGTAGAAAAT |
| L[16][6] | TTGGGGTCGAGGGTGGCAACATATAAAAGAAAATCAAGTTTT |
| L[17][0] | CGTTTACCAGAGCAAAATCCCTTATAAATCAATCATAACCCT |
| L[17][1] | CGAGATAGGGTGGCATAGTAAGAGCAACACTAAAGAATAGCC |
| L[17][2] | AGGAATTACGATGAGTGTTGTTCCAGTTTGGATAACGCCAAA |
| L[17][3] | ACTATTAAAGACATTCAACTAATGCAGATACAACAAGAGTCC |
| L[17][4] | TAGGAATACCAACGTGGACTCCAACGTCAAAGAGTTGAGATT |
| L[17][5] | CGTCTATCAGGTTACAGGTAGAAAGATTCATCGGCGAAAAAC |
| L[17][6] | GAACAACATTAGCGATGGCCCACTACGTGAACCGAACTAACG |
| Ef | CATACAGGCAAACTCCAGCCAGCTTTCCGGCATCCAATAAAT |
| Jjf[0] | TAGCAAGCAAAAAACCAATCAATAATCGGCTGCCGCGCCCAA |
| Jjf[1] | TCATTCCAAGATTTTCATCGTAGGAATCATTATCTTTCCTTA |
| Jjf[2] | AAACAATTCGCAACTAATAGATTAGAGCCGTTTAGACTTTAC |
| Jjf[3] | GGAGCACTAAACAACTCGTATTAAATCCTTTAAATATCTTTA |
| Jjf[4] | ATTAATTTTAGGAATTGAGGAAGGTTATCTAGCCCGAACGTT |
| Jjs | GCCGTTTTTAACGGGTATTA |
| Jc[0][0] | ACCCTCAGAGCTCTGGTGCCGGAAACCAGGCACAGAACCGCCCTATTAAA |
| Jc[0][1] | CGCCATTCAGGCACCCTCAGAACCGCCACCCTAAGCGCCATTCCGACTTG |
| Jc[0][2] | TTTAGTACCGCCTGCGCAACTGTTGGGAAGGGACTCAGGAGGGCTGCACC |
| Jc[0][3] | GGGCCTCTTCGCCGGAATAGGTGTATCACCGTCGATCGGTGCCTGCGAAG |
| Jc[0][4] | ATAAGTATAGCCTATTACGCCAGCTGGCGAAAGAGGGTTGATACTAAGCT |
| Jc[0][5] | CTGCAAGGCGACAGGCGGATAAGTGCCGTCGAGGGGGATGTGCCATCCAG |
| Jc[0][6] | TTGCTCAGTACTTAAGTTGGGTAACGCCAGGGTAGCGGGGTTCACTGAAG |
| Jc[1][0] | GCGTCATACATATTTGCACGTAAAACAGAAATGTTCCAGTAATAACAGCC |
| Jc[1][1] | CGTAGATTTTCAGCGCAGTCTCTGAATTTACCAAAGAAATTGTGGATAGC |
| Jc[1][2] | CCAGAATGGAAAGGTTTAACGTCAGATGAATACTCATTAAAGTACGGAGC |
| Jc[1][3] | GTACCTTTTACATATTCACAAACAAATAAATCTACAGTAACACTTCTGGA |
| Jc[1][4] | GATTGGCCTTGATCGGGAGAAACAATAACGGACAGGTCAGACCAGACCGA |
| Jc[1][5] | TGCTTTGAATAAGCATTGACAGGAGGTTGAGGTTCGCCTGATGTCTAAGA |
| Jc[1][6] | GAGCCGCCGCCCCAAGTTACAAAATCGCGCAGACCACCACCATCCAGAAG |
| Jcjm[0] | ACCGCACTTGAGGATTTAGAAGTACAATAGATAGCCAGAT |
| Jcjm[1] | AATACATTCATCGAGAACAAGCAAAACCAAGTATATAGGC |

| | |
|---|---|
| Jo[0][0] | ACCCTCAGAGCTAGCGGGGTTTTGCTCAGTACCAGAACCGCCCCATACGT |
| Jo[0][1] | TAACGCCAGGGTCTGGTGCCGGAAACCAGGCATTAAGTTGGGAATATTCT |
| Jo[1][0] | GCGTCATACATACCACCACCAGAGCCGCCGCCGTTCCAGTAACGGCATCA |
| Jo[1][1] | AAATCGCGCAGATTTGCACGTAAAACAGAAATCCAAGTTACAATGTTCTT |

In the reconfiguration process, releaser strands with toeholds are used for all the Jc, Jcjm and Jo. They base-pair with the target staples completely. The name of a releaser strand starts with 'r' in front of its target staple.

| Name | Sequence (5' to 3') of Releaser Strand |
|---|---|
| rJc[0][0] | TTTAATAGGGCGGTTCTGTGCCTGGTTTCCGGCACCAGAGCTCTGAGGGT |
| rJc[0][1] | CAAGTCGGAATGGCGCTTAGGGTGGCGGTTCTGAGGGTGCCTGAATGGCG |
| rJc[0][2] | GGTGCAGCCCTCCTGAGTCCCTTCCCAACAGTTGCGCAGGCGGTACTAAA |
| rJc[0][3] | CTTCGCAGGCACCGATCGACGGTGATACACCTATTCCGGCGAAGAGGCCC |
| rJc[0][4] | AGCTTAGTATCAACCCTCTTTCGCCAGCTGGCGTAATAGGCTATACTTAT |
| rJc[0][5] | CTGGATGGCACATCCCCCTCGACGGCACTTATCCGCCTGTCGCCTTGCAG |
| rJc[0][6] | CTTCAGTGAACCCCGCTACCCTGGCGTTACCCAACTTAAGTACTGAGCAA |
| rJc[1][0] | GGCTGTTATTACTGGAACATTTCTGTTTTACGTGCAAATATGTATGACGC |
| rJc[1][1] | GCTATCCACAATTTCTTTGGTAAATTCAGAGACTGCGCTGAAAATCTACG |
| rJc[1][2] | GCTCCGTACTTTAATGAGTATTCATCTGACGTTAAACCTTTCCATTCTGG |
| rJc[1][3] | TCCAGAAGTGTTACTGTAGATTTATTTGTTTGTGAATATGTAAAAGGTAC |
| rJc[1][4] | TCGGTCTGGTCTGACCTGTCCGTTATTGTTTCTCCCGATCAAGGCCAATC |
| rJc[1][5] | TCTTAGACATCAGGCGAACCTCAACCTCCTGTCAATGCTTATTCAAAGCA |
| rJc[1][6] | CTTCTGGATGGTGGTGGTCTGCGCGATTTTGTAACTTGGGGCGGCGGCTC |
| rJcjm[0] | ATCTGGCTATCTATTGTACTTCTAAATCCTCAAGTGCGGT |
| rJcjm[1] | GCCTATATACTTGGTTTTGCTTGTTCTCGATGAATGTATT |
| rJo[0][0] | ACGTATGGGGCGGTTCTGGTACTGAGCAAAACCCCGCTAGCTCTGAGGGT |
| rJo[0][1] | AGAATATTCCCAACTTAATGCCTGGTTTCCGGCACCAGACCCTGGCGTTA |
| rJo[1][0] | TGATGCCGTTACTGGAACGGCGGCGGCTCTGGTGGTGGTATGTATGACGC |
| rJo[1][1] | AAGAACATTGTAACTTGGATTTCTGTTTTACGTGCAAATCTGCGCGATTT |

### S6. References


1 Douglas, S. M., Marblestone, A. H., Teerapittayanon, S., Vazquez, A., Church, G. M. & Shih, W. M. Rapid Prototyping of 3D DNA-Origami Shapes with CaDNAno. *Nucleic Acids Research* **37**, 5001-5006 (2009).
